# *Phyllobacterium meliloti* sp. nov. a novel non-symbiotic bacterium isolated from root nodules of *Melilotus albus* (white sweet clover) grown in Canada

**DOI:** 10.1101/2024.11.22.624894

**Authors:** Eden S. P. Bromfield, Sylvie Cloutier, Michael F. Hynes

## Abstract

Two novel bacterial strains isolated from root-nodules of white sweet clover (*Melilotus albus*) plants grown at a Canadian site were previously characterized and placed in the genus *Phyllobacterium*. Here we present phylogenomic and phenotypic data to support the description of strain T1293^T^ as representative of a novel species and present the first complete closed genome sequence of a bacterial strain (T1018) representing the species ‘*P. pellucidum’*.

Phylogenetic analysis of genome sequences as well as analysis of 53 core genes placed novel strain T1293^T^ in a highly supported cluster of strains distinct from named *Phyllobacterium* species with *P. myrsinacearum* and *P. calauticae* as closest relatives. The highest average nucleotide identity (ANI) and digital DNA-DNA hybridization (dDDH) values of genome sequences of T1293^T^ compared to closest species type strains (84.1% and 26.5%, respectively) are well below the threshold values for bacterial species circumscription. The genome of strain T1293^T^ has a size of 5074034 bp with a DNA G+C content of 55 mol% and possesses three plasmids with sizes of 397619 bp, 476847 bp and 519835 bp. Detected in the genome were Type III and Type VI secretion system genes, implicated in plant-microbe and microbe-microbe interactions, but key nodulation, nitrogen-fixation and photosystem genes were not detected. Further analysis revealed that T1293^T^, like other *Phyllobacterium* species, possesses key genes encoding an enzyme complex implicated in the degradation of glyphosate, a widely used broad-spectrum herbicide that has negative consequences for many microorganisms including the human gut microbiome. A novel prophage (size ∼ 41.5 kb) was also detected in the genome of T1293^T^. Data for multiple phenotypic tests complemented the sequence-based characterization of strain T1293^T^. The data presented support the description of a new species and the name *Phyllobacterium meliloti* sp. nov. is proposed with T1293^T^ = LMG32641^T^ = HAMBI 3765^T^ as the species type strain.

## Introduction

The bacterial genus *Phyllobacterium*, placed in the family *Bartonellaceae* [1], was originally named after bacteria isolated from leaf nodules of tropical ornamental plants [2]. The 17 *Phyllobacterium* species described to date [3] are either free living or associated with plants; only two species (*P. sophorae* [4] and *P. trifolii* [5]) are able to form a symbiotic association with a plant host.

In a previous study [6], bacteria were isolated from root-nodules of *Melilotus albus* (white sweet clover) grown at a field site in Canada that had no history of cultivation. Phylogenetic analysis of four core gene sequences placed these bacterial isolates in several distinct lineages that were assigned to the genera *Ensifer*, *Rhizobium* and *Phyllobacterium* [7]. Two of these lineages, represented by strains T1293^T^ and T1018, were placed in the genus *Phyllobacterium* and shown to be novel genospecies. Since that time, a strain designated BT25^T^, closely related to strain T1018, was isolated from soil in Korea and described as a species named ‘*Phyllobacterium pellucidum*’ [8].

In the present work, we carried out detailed phylogenetic, genomic and phenotypic analyses of novel strain T1293^T^ and sequence based analyses of strain T1018. Based on the data generated we propose a novel species named *P. meliloti* sp. nov with T1293^T^ as the type strain and present the first complete closed genome sequence of a bacterial strain (T1018) representing the species ‘*P. pellucidum’*.

### Novel bacteria, habitat and isolation

Novel bacterial strains, *P. meliloti* sp. nov T1293^T^ and ‘*P. pellucidum’* T1018 were isolated in 1992 from root-nodules of *Melilotus albus* cultivar Polara plants grown at an uncultivated field site in Ottawa, Ontario, Canada that had no history of agriculture; the soil at this site was a sandy loam (pH 6.1, water) with good drainage [6].

*P. meliloti* sp. nov T1293^T^ was deposited in the BCCM/LMG Bacteria Collection, University of Ghent, Belgium as collection no. LMG32641^T^ and in the HAMBI Microbial Culture Collection, University of Helsinki, Finland as collection no. HAMBI 3765^T^.

Novel strain, ‘*P. pellucidum’* T1018 was deposited in the BCCM/LMG Bacteria Collection, University of Ghent, Belgium as LMG collection no. LMG 32375.

### DNA sequencing, genomic and phylogenetic analyses

Genomic DNAs were extracted and purified from bacterial cells grown for 7 days at 28 °C on Yeast-extract mannitol (YEM) agar medium as previously described [9].

The complete genomes of novel strains T1293^T^ and T1018 were sequenced at the Genome Quebec Innovation Centre, Canada, using Pacific Biosciences (PacBio) Sequel Single-Molecule Real-Time (SMRT) technology as detailed previously [9]; Flye software (version 2.9) [10] was employed for sequence assembly.

Estimated genome coverage for novel strain T1293^T^ was 3369-fold with 91069 polymerase reads and an average read length of 123841 bp; for strain T1018, coverage was 2247-fold with 61355 polymerase reads and an average read length of 93837 bp. The complete closed genome sequences of strains T1293^T^ and T1018 were of high quality [11] with completeness values of 99.0% (100^th^ percentile) and 99.4% (100^th^ percentile) and low contamination values of 1.2% and 0.9%, respectively.

The complete closed genome sequence of novel strain T1293^T^ has a size of 5074034 bp with DNA G+C content of 55 mol% (Table 1). Three circular plasmids possessing *repABC* genes encoding proteins involved in plasmid replication and segregation [12], were detected in this novel strain as follows: pT1293a (397619 bp), pT1293b (476847 bp) and pT1293c (519835 bp). It is noteworthy, that the detection of three plasmids in the genome of strain T1293^T^ is supported by the results of plasmid profile analysis (agarose gel electrophoresis) in our previous work [7].

**Table 1.**
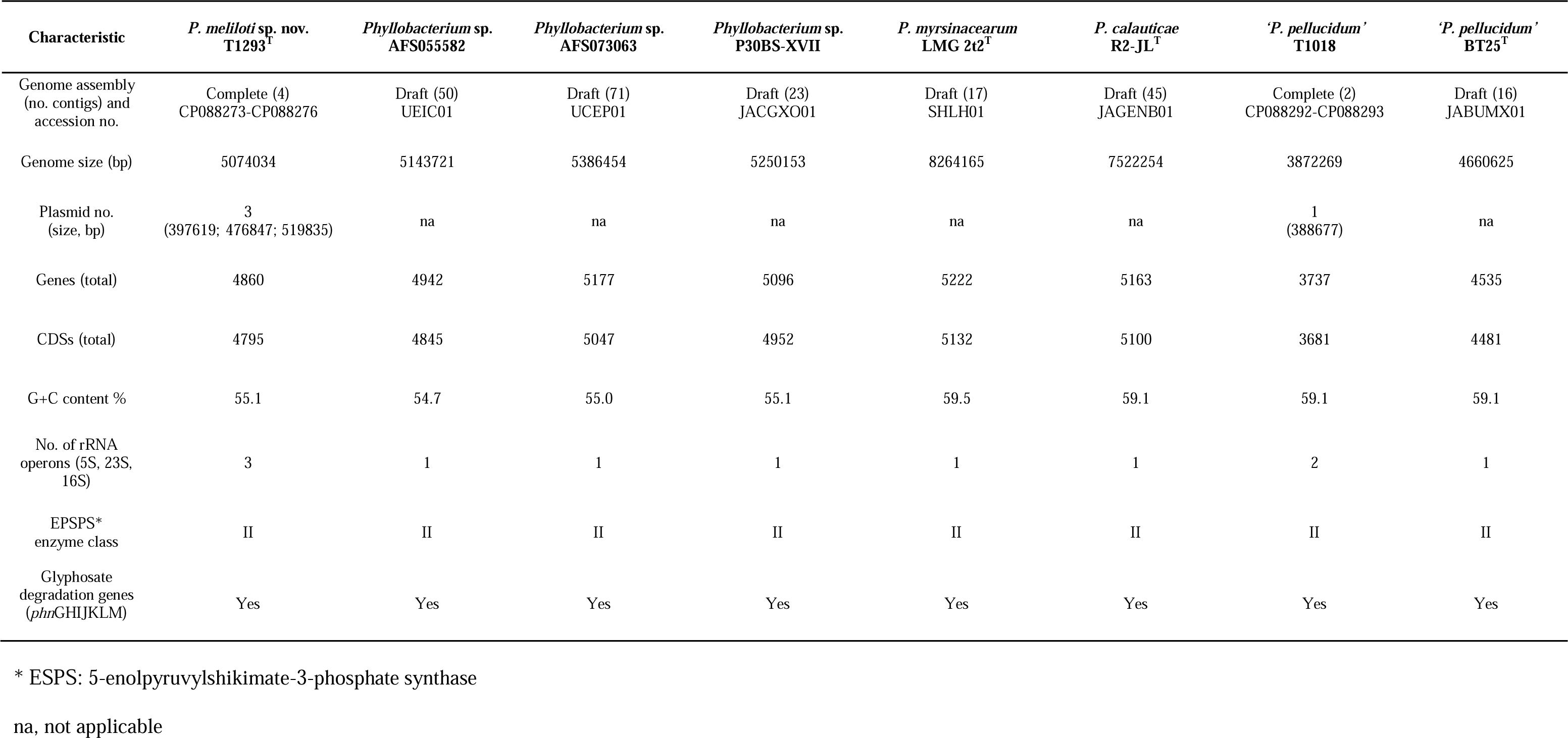
Characteristics of genome sequences of *Phyllobacterium meliloti* sp. nov. T1293^T^, ‘*Phyllobacterium pelludicum*’ T1018 and reference strains.

| Characteristic | *P. meliloti* sp. nov.<br>T1293<sup>T</sup> | *Phyllobacterium* sp.<br>AFS055582 | *Phyllobacterium* sp.<br>AFS073063 | *Phyllobacterium* sp.<br>P30BS-XVII | *P. myrsinacearum*<br>LMG 2t2<sup>T</sup> | *P. calauicae*<br>R2-JL<sup>T</sup> | ‘*P. pelludicum*’<br>T1018 | ‘*P. pelludicum*’<br>BT25<sup>T</sup> |
| --- | --- | --- | --- | --- | --- | --- | --- | --- |
| Genome assembly<br>(no. contigs) and<br>accession no. | Complete (4)<br>CP088273-CP088276 | Draft (50)<br>UEIC01 | Draft (71)<br>UCEP01 | Draft (23)<br>JACGXO01 | Draft (17)<br>SHLH01 | Draft (45)<br>JAGENB01 | Complete (2)<br>CP088292-CP088293 | Draft (16)<br>JABUMX01 |
| Genome size (bp) | 5074034 | 5143721 | 5386454 | 5250153 | 8264165 | 7522254 | 3872269 | 4660625 |
| Plasmid no.<br>(size, bp) | 3<br>(397619; 476847; 519835) | na | na | na | na | na | 1<br>(388677) | na |
| Genes (total) | 4860 | 4942 | 5177 | 5096 | 5222 | 5163 | 3737 | 4535 |
| CDSs (total) | 4795 | 4845 | 5047 | 4952 | 5132 | 5100 | 3681 | 4481 |
| G+C content % | 55.1 | 54.7 | 55.0 | 55.1 | 59.5 | 59.1 | 59.1 | 59.1 |
| No. of rRNA<br>operons (5S, 23S,<br>16S) | 3 | 1 | 1 | 1 | 1 | 1 | 2 | 1 |
| EPSPS*<br>enzyme class | II | II | II | II | II | II | II | II |
| Glyphosate<br>degradation genes<br>(*phn*GHJKLM) | Yes | Yes | Yes | Yes | Yes | Yes | Yes | Yes |

\* ESPS: 5-enolpyruvylshikimate-3-phosphate synthase
na, not applicable

The complete genome sequence of ‘*P. pellucidum’* T1018 has a size of 3872269 bp with DNA G+C content of 59 mol%. A single plasmid (pT1018 with size 388677bp) possessing *repABC* genes was detected in the genome of this strain (Table 1).

For phylogenetic analyses, nucleotide sequences were retrieved from genome sequences and alignments performed using MUSCLE [13]. Phylogenetic analyses were done using MrBayes (software version 3.2.1) with default priors as described previously [14]. Best fit substitution models were selected using ModelTest-NG [15] implemented in the web-based CIPRES Science Gateway version 3.3 [16]. In a previous study [17], we reported that the *recA* gene (encoding recombinase A involved in DNA repair) is a useful phylogenetic marker for rapid screening of bacterial isolates for potential novel genospecies. We performed BLASTn searches using the complete *recA* gene sequence of novel strain T1293^T^ (locus tag, LLE53_003560) as query to locate sequences of any closely related strains in NCBI databases. Based on the results of these searches we included the following unclassified but closely related strains (≥ 98.9% identity) to T1293^T^ in our phylogenetic and genomic analyses: AFS055582 and AFS073063 isolated respectively from an insect and a root of a *Zea mays* (corn) plant [18], and, strain P30BS-XVII isolated from pastureland bulk soil [19]

In accordance with minimal standards proposed for the use of genome data for prokaryotic taxonomy [20], we verified that the 16S rRNA gene sequences of strains T1293^T^ and T1018 generated by the Sanger method [14] were identical to the respective 16S rRNA sequences generated by whole genome sequencing.

The 16S rRNA gene is highly conserved in bacteria, and as such, its usefulness as a taxonomic tool is limited to assessment of genus level or higher taxonomic rank [20]. A Bayesian phylogenetic tree (Fig. S1) of nearly full length 16S rRNA gene sequences (alignment length 1364 positions) of type strains of all 17 described *Phyllobacterium* species confirms placement of novel strains T1293^T^ and T1018 in the genus *Phyllobacterium.* Strain T1293 ^T^ is placed in a distinct cluster with the unclassified strains AFS055582, AFS073063 and P30BS-XVII that were retrieved from database searches. Placement in the 16S rRNA gene tree also indicates that closest species relatives of T1293^T^ are type strains of *P. myrsinacearum* and *P. calauticae* while the closest relative of T1018 is the type strain of ‘*P. pellucidum’*.

To further verify the taxonomic position of strains T1293^T^ and T1018 we carried out the following phylogenetic analyses employing strains AFS055582, AFS073063, P30BS-XVII and 15 type strains of *Phyllobacterium* species as reference taxa: 1) multiple locus sequence analysis (MLSA) of 53 concatenated single-copy core gene sequences encoding ribosome protein subunits (*rps*) [21], and, 2) analysis of whole genome sequences implemented in the Type Strain Genome Server (TYGS) [22]. Procedural details of these phylogenetic analyses were as described previously [23].

The topology of the Bayesian phylogenetic tree based on 53 core gene sequences (Fig. 1) and the topology of the tree based on genome sequences (Fig. S2) confirm that T1293^T^ is placed with strains AFS055582, AFS073063, P30BS-XVII in a highly supported cluster distinct from named *Phyllobacterium* species with the type strains of *P. myrsinacearum* and *P. calauticae* as closest relatives. The two trees further show that novel strain T1018 is placed in a highly supported lineage together with the type strain of *‘P. pellucidum’* as closest relative.

**Figure 1.**
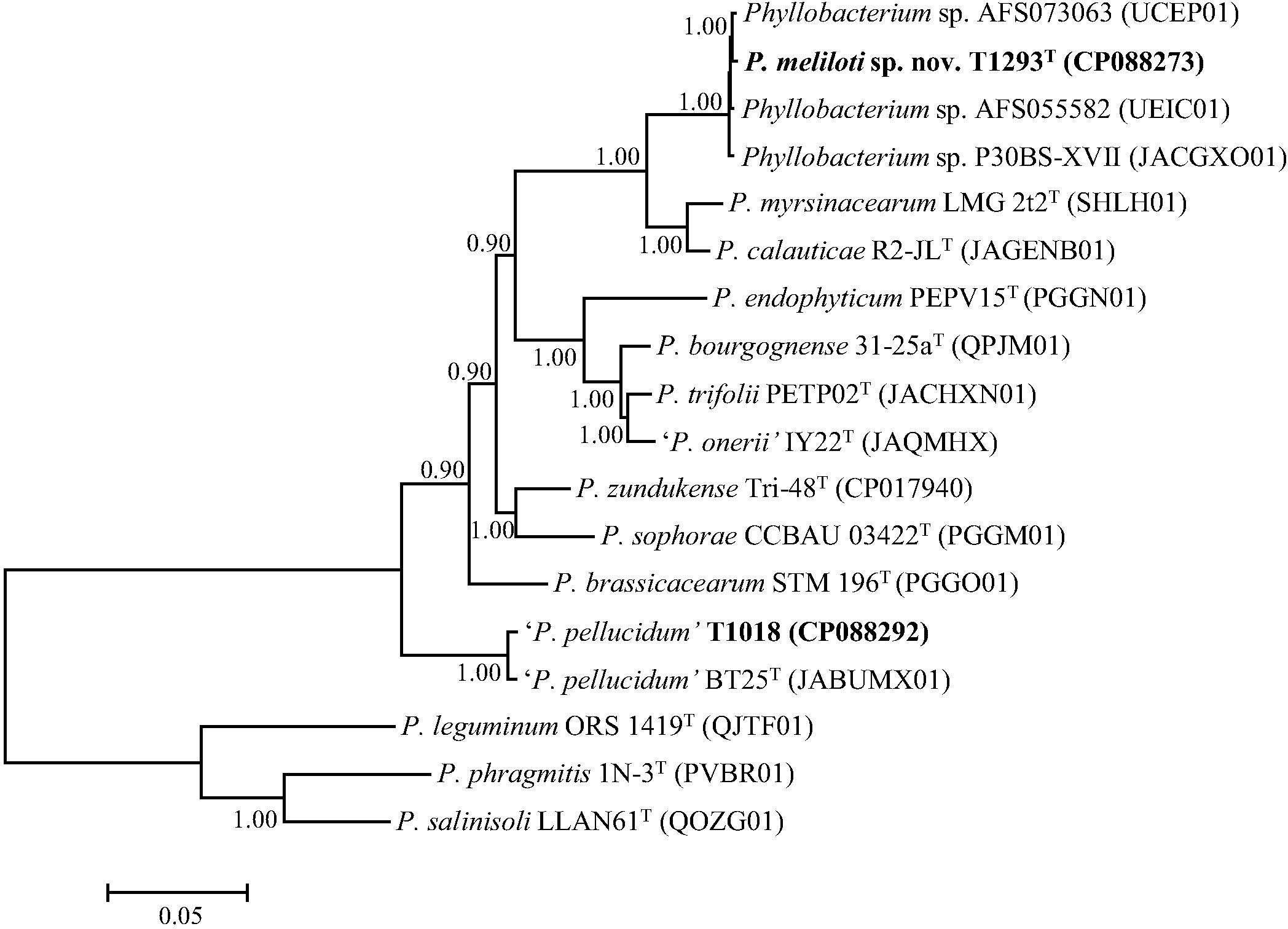
Bayesian phylogenetic tree (GTR+G+I substitution model) inferred from 53 full-length concatenated ribosome protein subunit (*rps*) gene sequences for *Phyllobacterium meliloti* sp. nov. strain T1293^T^ and reference taxa. Alignment length, 22868 positions. NCBI sequence accession numbers are given in parentheses. Posterior probabilities ≥ 0.90 are shown. Bar, expected substitutions per site.

Average nucleotide identity (ANI) and digital DNA–DNA hybridization (dDDH) were employed as indices of overall genomic relatedness [20] to facilitate bacterial species delineation. ANI values were calculated using FastANI [24] and the established ANI threshold of 95–96% was employed for species circumscription [20]. Overall genomic relatedness values based on dDDH were calculated using algorithms implemented in the web-based Type Strain Genome Server (TYGS) and the established dDDH threshold of 70% was used to delineate species boundaries [22]. ANI and dDDH values for pair-wise comparisons of the genome sequence of T1293^T^ with genome sequences of reference strains (AFS055582, AFS073063, P30BS-XVII and species type strains of closest relatives) are presented in Table 2. The highest ANI and dDDH values were for comparisons of T1293^T^ with type strains of *P. calauticae* (ANI, 84.4%; dDDH, 27%) and *P. myrsinacearum* (ANI, 84.1%; dDDH, 26.5 %). These values are well below the accepted thresholds for species delineation and confirm that T1293^T^ represents a novel species within the genus *Phyllobacterium*. Values for genome sequences of strains AFS055582, AFS073063, P30BS-XVII compared with T1293^T^ were above the respective cutoff values of 95-96% (ANI) and 70% (dDDH) for species circumscription, confirming that these strains belong to the same species cluster as T1293^T^.

**Table 2.**
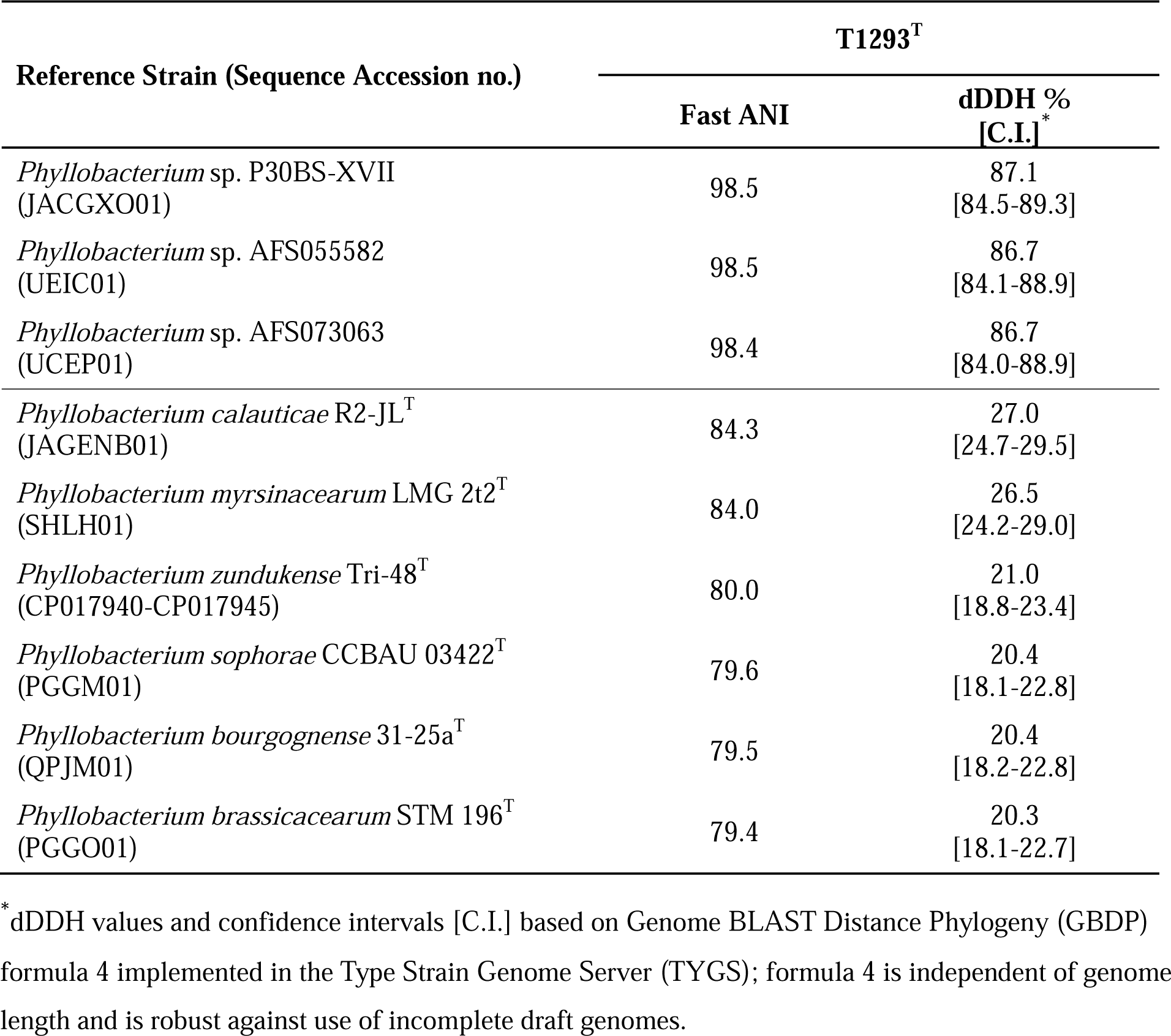
Average Nucleotide Identity (ANI) and digital DNA–DNA hybridization (dDDH) values for pair-wise comparisons of genome sequences of *Phyllobacterium meliloti* sp. nov. T1293^T^ (Accession no. CP088273-CP088276) with unclassified strains P30BS-XVII, AFS055582, AFS073063 (retrieved from NCBI database searches) and species type strains of closest relatives.

ANI and dDDH values for the comparison of T1018 with the closely related type strain of *‘P. pellucidum’* (98.2 % and 84.1%) were above the respective cutoff values of 95-96% and 70% for species circumscription indicating that novel strain T1018 is a member of the species ‘*P. pellucidum’*.

Further genome analyses were carried out using Geneious Prime 2023.0.4 software (www.geneious.com). Based on these analyses, key type III secretion system (T3SS) and type VI secretion system (T6SS) genes, implicated in plant-microbe and microbe-microbe interactions [25, 26, 27], were detected respectively in plasmids pT1293c and pT1293b of novel strain T1293^T^. Both of these secretion system gene clusters appear to be complete, but there are no obvious clues as to their function or what effector molecules they might export into other cells. Both of these gene clusters are most similar to those in other species of *Phyllobacterium*.

In contrast, key T3SS and T6SS genes were not detected in the genome of ‘*P. pellucidum’* strain T1018.

We did not detect key type II and IV secretion system (T2SS and T4SS) genes [28, 29], photosystem genes or genes involved in symbiosis (e.g., nodulation (*nod*) and nitrogen fixation (*nif*) genes), in the genomes of strains T1293^T^ and T1018. The absence of T4SSs on plasmids in the two novel strains indicates that these plasmids are not self-transmissible.

Genes in bacteria encoding 16S, 23S, and 5S rRNAs are linked together and expressed as a single ribosomal RNA operon (*rrn*) [30]. Many bacterial species possess multiple *rrn* operon copies in their genomes and these copies may be located either on chromosomes, plasmids or both [23, 30, 31, 32]. Our data for the genome analysis of novel strains T1293^T^ and T1018 (Table 1) are in line with these reports: strain T1293^T^ was found to possess three *rrn* copies with two copies located on the chromosome and one copy on plasmid, pT1293a (size, 397.619 kb), whereas, *‘P. pellucidum’* T1018 was found to possess two *rrn* copies, both on the chromosome. While we did not detect any intragenomic variation between multiple *rrn* operon copies in novel strains T1293^T^ or T1018, this type of heterogeneity is not without precedent and has been reported occasionally in different bacterial species [23, 30, 33].

Infectious viruses of bacteria (called bacteriophages or phages) are mediators of horizontal gene transfer and as a consequence represent an important driver of bacterial diversification and evolution [33]. In this connection, we used web based software, PHASTEST [35] to detect potential prophage sequences within the genomes of novel strains T1293^T^ and T1018. Two potential intact (as defined by PHASTEST) prophages were detected in the chromosome of T1293^T^: the first located at co-ordinates 940,853-982,445bp (size, 41.5kb) and the second located at coordinates 1,899,457-1,915,246bp (size 15.7kb). The 41.5 kb predicted prophage did not have any close homologues at the nucleotide level, with the best BlastN hit being a to the *Phyllobacterium* A18/5-2 chromosome (Accession Number CP104966.1; 80 % identity over a short region of 4916 nucleotides). This predicted prophage sequence contains an integrase gene, genes predicted to be involved in phage replication, and a complete set of predicted structural genes from phage assembly, so it is very likely to encode a functional phage, which represents a potential founding member of a novel genus within the class *Caudoviricetes* (International Committee on Taxonomy of Viruses (ICTV) (https://ictv.global/taxonomy/)). Individual genes encoded predicted proteins with highest similarity to a variety of phages from organisms like *Sinorhizobium meliloti*, *Paracoccus* species, and *Burkholderia* species, and the putative prophage sequence is upstream of several tRNA genes, with tRNAs often being the site of phage integration. The second predicted prophage region (15.7kb) encoded a complete set of structural proteins with several showing most similarity to proteins of *Rhodobacter* phage RcCronus (NCBI reference: NC_042049) and other *Rhodobacter* phages. However, due to the fact that the region is too small to encode a complete phage genome, and the absence of genes encoding integrases, DNA replication enzymes and other functions, it seems unlikely that this is a functional prophage. In addition, some genes encode proteins that are most similar to those from gene transfer agents (GTA) [36] of *Rhodobacter* spp. and relatives, raising the possibility that this DNA region encodes a GTA. A potential prophage similar to the 15.7 kb sequence was also detected in the chromosome of ‘*P. pellucidum*’ T1018 (co-ordinates 1,641,578-1,655,055bp; size, 13.4 kb), but due to its size and lack of some key genes might be a GTA-encoding region.

Glyphosate is the most extensively used non-specific herbicide in the world for food production [37]. Its unprecedented scale of use has resulted in negative consequences for human health, plants, fungi and bacteria including the human gut microbiome [37, 38, 39]. Consequently there is considerable interest in deploying bacteria that have potential to degrade glyphosate to facilitate the bioremediation of contaminated environments [19, 37].

Glyphosate specifically inhibits the enzyme, 5-enolpyruvylshikimate-3-phosphate synthase (EPSPS) found only in plants, fungi and bacteria and interferes with the synthesis of essential aromatic amino acids in the shikimate pathway [39, 40]. EPSPS enzymes have been placed in two major classes based on their sensitivity to glyphosate and their sequence variations: Class I enzymes (found in plants and most Gram-negative bacteria, such as *E.coli*) are naturally sensitive to glyphosate whereas Class II enzymes share less than 30% sequence similarity with class I enzymes, retain their efficiency even in the presence of elevated glyphosate concentrations and are found in some naturally occurring glyphosate-tolerant bacteria such as strains of *Agrobacterium* [19, 41, 42, 43].

Massot et al [19] reported that unclassified *Phyllobacterium* sp. strain P30BS-XVII was highly resistant to glyphosate with a minimal inhibitory concentration (MIC) of 10,000 mg glyphosate/ kg. It should be noted that in the present work we show that strain P30BS-XVII belongs to the same species cluster as novel strain T1293^T^ (see Fig. 1 and Table 2).

We analyzed the amino acid sequences of EPSP enzymes in novel strains T1293^T^ and ‘*P. pelluducum’* T1018 as well as reference taxa consisting of strains P30BS-XVII, AFS055582, AFS073063, *P. myrsinacearum* LMG 2t2^T^, *P. calauticae* R2-JL^T^ and *P. pellucidum* BT25^T^ using software implemented in the *EPSPSClass* web server [39]; the results revealed that novel strains and all reference strains possess Class II glyphosate tolerant enzymes.

Further genomic analysis showed that T1293^T^, T1018 and all reference strains in Table 1 also possess key genes (*phn*GHIJKLM) encoding the C-P lyase enzyme complex necessary for the degradation of glyphosate [43]. Collectively these data suggest that glyphosate tolerance is a common feature of *Phyllobacterium* spp. As such, strains T1293^T^ and T1018 as well as other strains of *Phyllobacterium* spp. identified in this work constitute a potential new resource for furthering research on the bioremediation of glyphosate contaminated environments.

### Phenotypic characterization

Colonies of strain T1293^T^ are translucent, beige coloured, spreading with copious gum with diameters of about 1-2 mm after growth on YEM agar medium for 3 days at 28 °C. Cells of T1293^T^ are Gram-stain-negative based on the method of Buck [44]. Grows in the presence of 2% NaCl, at pH5 and pH10 and at temperatures of 10°C and 37°C after 2 days on YEM agar medium (Table S1).

Cell morphology was investigated using cells grown in YEM broth for two days at 28 °C; cells were visualized using transmission (model, H-7000; Hitachi) and scanning (model, Hitachi SU7000 FESEM) electron microscopes as described previously [23]. The results revealed that cells of T1293^T^ are rod shaped, and possess one or more flagella (Fig. S3).

Further phenotypic characterization using API^®^ strips (API 20NE) and BIOLOG GENIII MicroPlates™ was done at BCCM/LMG, Ghent University, Belgium. The results (Table S2), show that novel strain T1293^T^ can be readily differentiated from close relatives *P. calauticae* R2-JL^T^ and *P. myrsinacearum* LMG 2t2^T^ as well as from *‘P. pellucidum’* T1018 and ‘*P. pellucidum’* BT25^T^, based on multiple phenotypic tests.

In our previous study, [7], novel strain T1293^T^ was found to be highly resistant to multiple antibiotics including carbenicillin (>1000 μg ml^−1^), kanamycin (>100 μg ml^−1^), erythromycin (∼100 μg ml^−1^) and tetracycline (>50 μg ml^−1^). These results were supported by our detection of multiple antibiotic resistance genes in the genome of strain T1293^T^ using software implemented in the BV-BRC web-based platform [45]. Antibiotic resistance genes detected included genes encoding enzymes that inactivate beta-lactam antibiotics (e.g., carbenicillin), macrolide antibiotics (e.g., erythromycin), amino-glycoside antibiotics (e.g., kanamycin and neomycin) as well as genes conferring resistance to tetracycline antibiotics (data not shown).

Analysis of fatty acids was carried out using the Sherlock Microbial Identification System (midi) version 6.0 and the RTSBA6 database as described previously [23]. The most abundant fatty acids in strain T1293^T^ were C16:0 (8.3%), C18:1ω7c 11-methyl (14.2%), C19:0 cyclo ω8c (46.8%) and C18:1ω6c/18:1ω7c (summed feature 8) (8.1%), consistent with the profiles of reference strains (Table S3) and with members of the genus *Phyllobacterium* reported previously [46].

Plant tests using modified Leonard jars were carried out as described previously [7]. Tests done in this study as well as in the previous study [7] showed that novel strain T1293^T^ did not elicit nodules on the roots of plants of *Melilotus albus* cv. Polara, *Medicago lupulina* (Black medic), *Medicago sativa* (alfalfa) or *Macroptilium atropurpureum* (Siratro).

### Description of Phyllobacterium meliloti sp. nov

*Phyllobacterium meliloti (me.li.lo’ti.* N.L. gen. n *meliloti,* of the plant genus *Melilotus)*.

Colonies are translucent, beige colored, spreading with copious gum formation and with diameters of ∼1-2 mm after growth on YEM agar medium for 3 days at 28 °C. Cells are Gram-stain-negative, aerobic, non-spore-forming rods with one or more flagella. Grows at pH 5 and pH10 (optimal at ∼ pH 6.8), at temperatures of 10°C and 37°C (optimal at ∼ 28°C) and in the presence of 2% NaCl, after 2 days on YEM agar medium.

Positive for the utilization of 40 carbon sources including D-raffinose, β-methyl-D glucoside, N-acetyl-β-D mannosamine, D-saccharic acid, D-galactose, citric acid and D-glucose-6-PO4. Does not utilize 15 carbon sources including: glucuronamide, methyl pyruvate, inosine, tween 40 and D-aspartic acid. Resistant to 12 chemical compounds including minocycline, niaproof 4 and potassium tellurite but susceptible to sodium bromate.

Positive for urease activity, esculin hydrolysis and assimilation of mannose, mannitol, N-acetyl-glucosamine, maltose, potassium gluconate, malate, trisodium citrate, and arabinose.

Negative for hydrolysis of gelatin, ß-galactosidase activity, arginine dihydrolase, reduction of nitrates to nitrites, reduction of nitrates to nitrogen, indol production and fermentation of glucose. Does not assimilate capric acid, phenylacetic acid, and adipic acid.

Predominant fatty acids (>10%) are C18:1ω7c 11-methyl and C19:0 cyclo ω8c.

The type strain, T1293^T^ (=LMG32641^T^ = HAMBI 3765^T^), was isolated from a root nodule of a *Melilotus albus* (white sweet clover) plant growing in Canada. The type strain possesses key type III secretion system (T3SS) and type VI secretion system (T6SS) genes, but does not possess key nodulation, nitrogen fixation or photosystem genes.

Does not elicit root-nodules on plants of *Melilotus albus* (white sweet clover), *Medicago lupulina* (black medic), *Medicago sativa* (alfalfa) or *Macroptilium atropurpureum* (Siratro).

The DNA G+C content of the type strain is 55.1% and the genome size is 5.07 Mbp. GenBank/EMBL/DDBJ accession numbers for the complete genome and the 16S rRNA gene sequence of the type strain are, respectively CP088273-CP088276 and EU928869.

## Acknowledgements

The authors are grateful to Keith Hubbard, Microscopy Centre, Agriculture and Agri-Food Canada, Ottawa for preparing electron microscope images.

## Author contributions

EB conceived and co-ordinated the project, received the funding, and wrote the draft manuscript. SC and EB carried out the experiments. SC, EB, and MH analyzed the data. All authors contributed to the article and approved the submitted version.

## Conflicts of interest

The authors declare that there are no conflicts of interest.

© His Majesty the King in Right of Canada as represented by the Minister of Agriculture and Agri-Food 2024 for the work done by Eden S.P. Bromfield and Sylvie Cloutier.

