## Supplementary Data for "*Phyllobacterium meliloti* sp. nov. a novel non-symbiotic bacterium isolated from root nodules of *Melilotus albus* (white sweet clover) grown in Canada"

**Supplementary Table S1.** Growth characteristics of *Phyllobacterium* strains: **1**. *P. meliloti* sp. nov. T1293$^T$, **2.** *P. myrsinacearum* LMG 2t2$^T$, **3**. *P. calauticae* R2-JL$^T$, **4.** '*P. pellucidum'* T1018, and, **5**. '*P. pellucidum'* BT25$^T$.

| **Characteristic** | **1** | **2** | **3** | **4** | **5** |
|---|---|---|---|---|---|
| Growth on YEM agar medium* | | | | | |
| 10 °C | ± | ± | ± | - | - |
| 37 °C | ± | ± | ± | ± | ± |
| pH 5 | + | + | + | + | + |
| pH 10 | + | + | + | + | ± |
| 0.5% NaCl | + | + | + | + | + |
| 1% NaCl | + | + | + | + | + |
| 2% NaCl | + | + | ± | - | ± |

* Positive, +; weak, ± ; negative, − ; not determined, ND. Values are based on three replicates after 48hrs.

**Supplementary Table S2.** Phenotypic characteristics of *Phyllobacterium* strains: **1**. *P. meliloti* sp. nov., T1293$^{T}$, **2.** *P. calauticae* R2-JL$^{T}$, **3.** *P. myrsinacearum* LMG 2t2$^{T}$, **4.** *'P. pellucidum'* T1018, and, **5**. '*P. pellucidum'* BT25$^{T}$.

| **Characteristic** | **1** | **2** | **3** | **4** | **5** | **Characteristic** | **1** | **2** | **3** | **4** | **5** |
|---|---|---|---|---|---|---|---|---|---|---|---|
| C-source utilization (Biolog) | | | | | | | | | | | |
| Dextrin | - | - | w | - | - | Glycyl-L-Proline | + | w | + | w | w |
| D-Maltose | + | + | + | - | - | L-Alanine | + | + | + | - | - |
| D-Trehalose | + | + | + | - | - | L-Arginine | w | w | + | w | w |
| D-Cellobiose | w | + | + | w | w | L-Aspartic Acid | + | + | + | + | + |
| Gentiobiose | + | + | + | w | w | L-Glutamic Acid | + | + | + | + | + |
| Sucrose | + | + | + | - | - | L-Histidine | + | w | + | + | + |
| D-Turanose | + | + | + | w | w | L-Pyroglutamic Acid | w | w | w | - | - |
| Stachyose | - | - | - | - | - | L-Serine | + | w | + | - | - |
| D-Raffinose | + | w | - | - | - | Pectin | w | w | + | - | - |
| α-D-Lactose | - | - | - | - | - | D-Galacturonic Acid | w | + | + | w | w |
| D-Melibiose | + | + | - | - | - | L-Galactonic Acid Lactone | w | + | + | w | w |
| β-Methyl-DGlucoside | + | w | w | - | - | D-Gluconic Acid | + | + | + | - | w |
| D-Salicin | - | - | - | - | - | D-Glucuronic Acid | w | + | + | + | w |
| N-Acetyl-DGlucosamine | + | + | + | + | + | Glucuronamide | - | + | + | w | w |
| N-Acetyl-β-DMannosamine | + | - | + | + | + | Mucic Acid | - | - | - | - | - |
| N-Acetyl-DGalactosamine | + | + | + | + | + | Quinic Acid | - | - | - | - | - |
| N-Acetyl Neuraminic Acid | - | - | + | - | - | D-Saccharic Acid | + | - | - | - | - |
| α-D-Glucose | w | + | + | w | w | p-HydroxyPhenylacetic Acid | - | - | - | - | - |
| D-Mannose | w | + | + | w | w | Methyl Pyruvate | - | w | + | - | - |
| D-Fructose | + | + | + | + | + | D-Lactic Acid Methyl Ester | w | w | + | w | w |
| D-Galactose | + | + | + | w | w | L-Lactic Acid | + | + | + | w | w |
| 3-Methyl Glucose | - | - | - | - | - | Citric Acid | + | + | + | - | - |
| D-Fucose | + | + | + | + | + | α-Keto-Glutaric Acid | w | w | + | w | w |
| L-Fucose | + | + | + | + | + | D-Malic Acid | + | + | + | + | + |
| L-Rhamnose | + | + | + | w | w | L-Malic Acid | + | + | + | + | + |
| Inosine | - | - | - | - | - | Bromo-Succinic Acid | + | w | + | + | + |
| D-Sorbitol | + | + | + | + | + | Tween 40 | - | - | w | - | - |
| D-Mannitol | + | + | + | + | + | γ-Amino-Butryric Acid | w | w | + | w | w |
| D-Arabitol | + | + | + | + | + | α-HydroxyButyric Acid | w | w | w | - | - |
| myo-Inositol | + | + | + | + | + | β-Hydroxy-D,LButyric Acid | + | + | + | + | + |
| Glycerol | + | + | + | w | w | α-Keto-Butyric Acid | + | w | w | + | + |
| D-Glucose- 6-PO4 | + | - | - | - | - | Acetoacetic Acid | w | w | w | w | w |
| D-Fructose- 6-PO4 | w | w | w | - | - | Propionic Acid | + | w | + | + | + |
| D-Aspartic Acid | - | - | - | - | - | Acetic Acid | + | + | + | + | + |
| Gelatin | - | - | - | - | - | Formic Acid | + | + | + | + | + |
| Chemical Sensitivity (Biolog) | | | | | | | | | | | |
| 1% Sodium Lactate | + | + | + | - | w | Vancomycin | + | + | + | w | w |
| Fusidic Acid | w | + | + | w | - | Tetrazolium Violet | + | + | + | + | + |
| D-Serine | w | w | - | - | - | Tetrazolium Blue | w | + | + | w | + |
| Troleandomycin | + | w | w | + | + | Nalidixic Acid | + | + | w | w | w |
| Rifamycin SV | + | + | + | + | + | Lithium Chloride | w | w | w | w | w |
| Minocycline | + | - | w | - | - | Potassium Tellurite | + | + | - | + | + |
| Lincomycin | + | + | + | + | + | Aztreonam | + | + | + | + | + |
| Guanidine HCl | w | w | w | - | w | Sodium Butyrate | + | w | w | w | w |
| Niaproof 4 | + | - | - | + | + | Sodium Bromate | - | w | w | - | - |
| Metabolism/Assimilation (API 20NE) | | | | | | | | | | | |
| Reduction of nitrates to nitrites | - | - | - | - | - | Assimilation of mannose | + | + | + | + | + |
| Reduction of nitrates to nitrogen | - | - | - | - | - | Assimilation of mannitol | + | + | + | + | + |
| Indol production | - | - | - | - | - | Assimilation of N-acetyl-glucosamine | + | + | + | + | + |
| Fermentation of glucose | - | - | - | - | - | Assimilation of maltose | + | + | + | - | - |
| Arginine dihydrolase | - | - | - | - | - | Assimilation of potassium gluconate | + | w | w | - | - |
| Urease activity | + | + | + | + | + | Assimilation of capric acid | - | - | - | - | - |
| Hydrolysis of esculin | + | + | + | w | w | Assimilation of adipic acid | - | - | - | - | - |
| Hydrolysis of gelatin | - | - | - | - | - | Assimilation of malate | + | + | + | + | + |
| ß-galactosidase activity | - | + | - | - | - | Assimilation of trisodium citrate | + | + | + | - | - |
| Assimilation of glucose | + | + | + | + | + | Assimilation of phenylacetic acid | - | - | - | - | - |
| Assimilation of arabinose | w | w | w | w | w | | | | | | |

BIOLOG GEN III MicroPlates and API 20NE (48h incubation at 28 ºC): +, Positive; w, weak; −, negative; v, variable. ND, not determined.

**Supplementary Table S3.** Fatty acid profiles of *Phyllobacterium* strains: **1**. *P. meliloti* sp. nov., T1293$^T$, **2.** *P. calauticae* R2-JL$^T$, **3.** *P. myrsinacearum* LMG 2t2$^T$, **4.** *'P. pellucidum'* T1018, and, **5**. '*P. pellucidum'* BT25$^T$.

| **Fatty Acid** | **1** | **2** | **3** | **4** | **5** |
|---|---|---|---|---|---|
| 13:1 at 12-13 | - | - | 0.2 | - | - |
| 14:0 | - | - | 0.2 | 0.4 | 0.62 |
| 15:0 3OH | 0.4 | - | 0.3 | - | - |
| 16:0 | 8.3 | 6.3 | 8.0 | 13.9 | 17.3 |
| 16:0 N alcohol | - | - | - | - | 0.6 |
| 16:0 2OH | 0.5 | - | 0.2 | - | - |
| 16:0 3OH | 7.1 | 5.0 | 5.6 | 0.8 | 1.1 |
| 16:1 ω11c | 0.4 | - | 0.4 | - | - |
| 17:0 cyclo | 1.8 | 1.7 | 2.2 | 1.8 | 2.6 |
| 17:0 3OH | 0.3 | - | 0.2 | - | - |
| 17:1 ω8c | 0.5 | - | 0.3 | 0.4 | - |
| 17:0 | 0.3 | - | 0.2 | 0.7 | 1.4 |
| 18:0 | 1.5 | 1.2 | 1.3 | 1.4 | 1.3 |
| 18:0 iso | 0.3 | - | 0.6 | - | - |
| 18:1 2OH | 4.9 | - | 2.0 | - | - |
| 18:0 2OH | 1.0 | - | - | - | - |
| 18:0 3OH | - | 1.5 | 1.6 | - | - |
| 18:1 ω9c | - | - | 0.7 | - | - |
| 18:1ω7c 11-methyl | 14.2 | 7.6 | 6.3 | 17.3 | 14.2 |
| 18:3 ω6c (6,9,12) | - | - | - | - | 1.6 |
| 19:0 | 0.6 | - | - | - | - |
| 19:0 cyclo ω8c | 46.8 | 63.8 | 56.9 | 35.5 | 32.5 |
| 20:0 | - | - | - | - | 0.8 |
| 20:1 ω7c | 0.4 | - | 0.4 | - | - |
| 20:2 ω6,9c | - | 2.2 | 1.2 | 1.5 | 3.2 |
| Summed feature 2* | 1.8 | 2.0 | 1.7 | 0.6 | 2.0 |
| Summed feature 3* | 0.3 | - | 0.5 | 1.8 | 1.4 |
| Summed feature 5* | - | 0.6 | - | 0.4 | 0.9 |
| Summed feature 7* | 0.5 | 1.0 | 0.2 | 1.7 | 1.5 |
| Summed feature 8* | 8.1 | 7.2 | 8.9 | 21.9 | 17.1 |

* Summed Features are fatty acids that cannot be resolved reliably from another fatty acid using the chromatographic conditions chosen. The MIDI system groups these fatty acids together as one feature with a single percentage of the total. Summed feature 2, 12:0 aldehyde/?; Summed feature 3, 16:1 ω6c/16:1ω7c; Summed feature 5, 18:0 ante/18:2 ω6,9c; Summed feature 7, 19:1 ω7c/19:1 ω6c; summed feature 8, 18:1ω6c/18:1ω7c;

-, not detected.

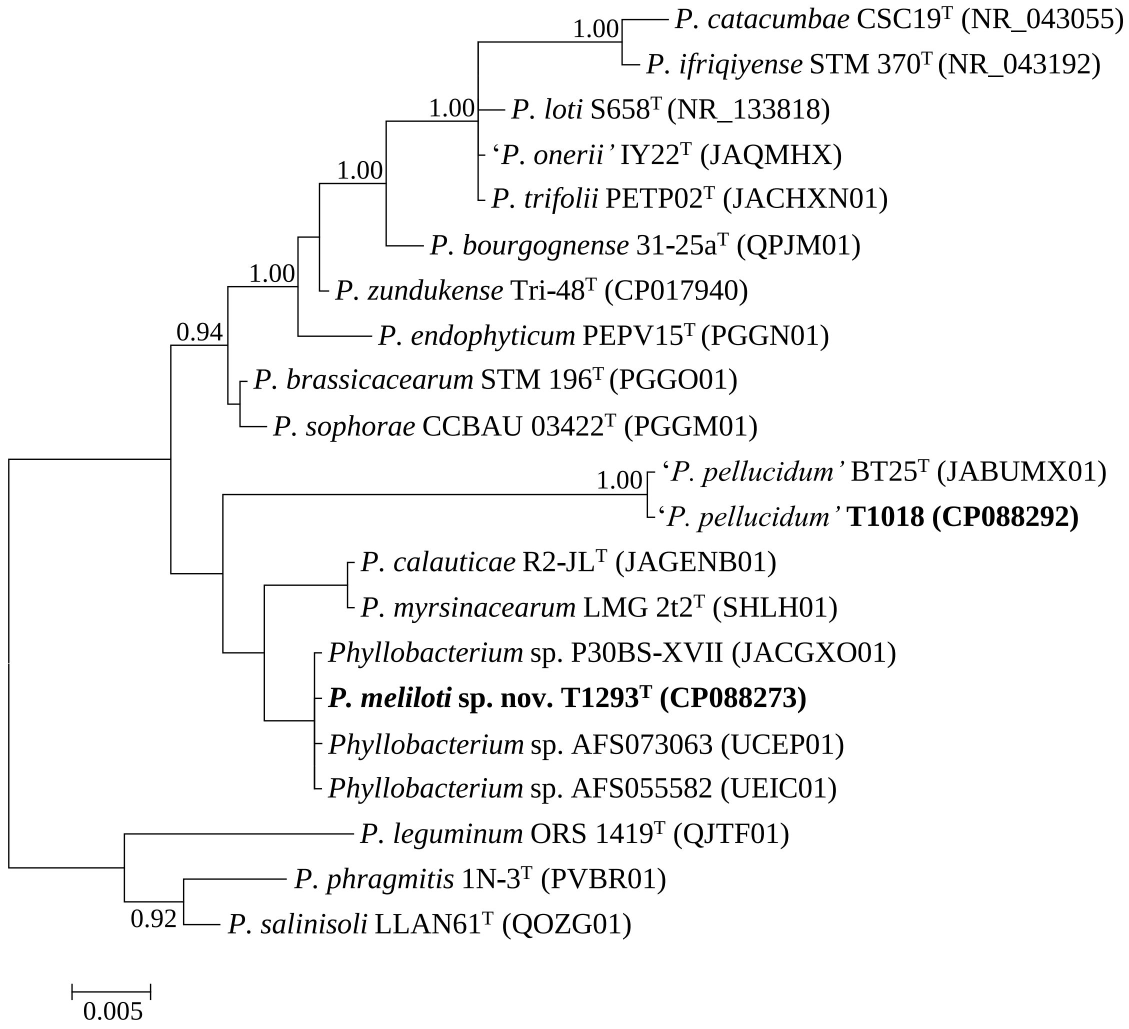


**Supplementary Figure S1**. Bayesian phylogenetic tree of 16S rRNA gene sequences (1364 positions) for *Phyllobacterium meliloti* sp. nov. strain T1293$^{T}$ and reference taxa (GTR + G + I substitution model). NCBI sequence accession numbers are given in parentheses. Only posterior probabilities ≥ 0.9 are shown. Scale bar represents expected number of substitutions per site.

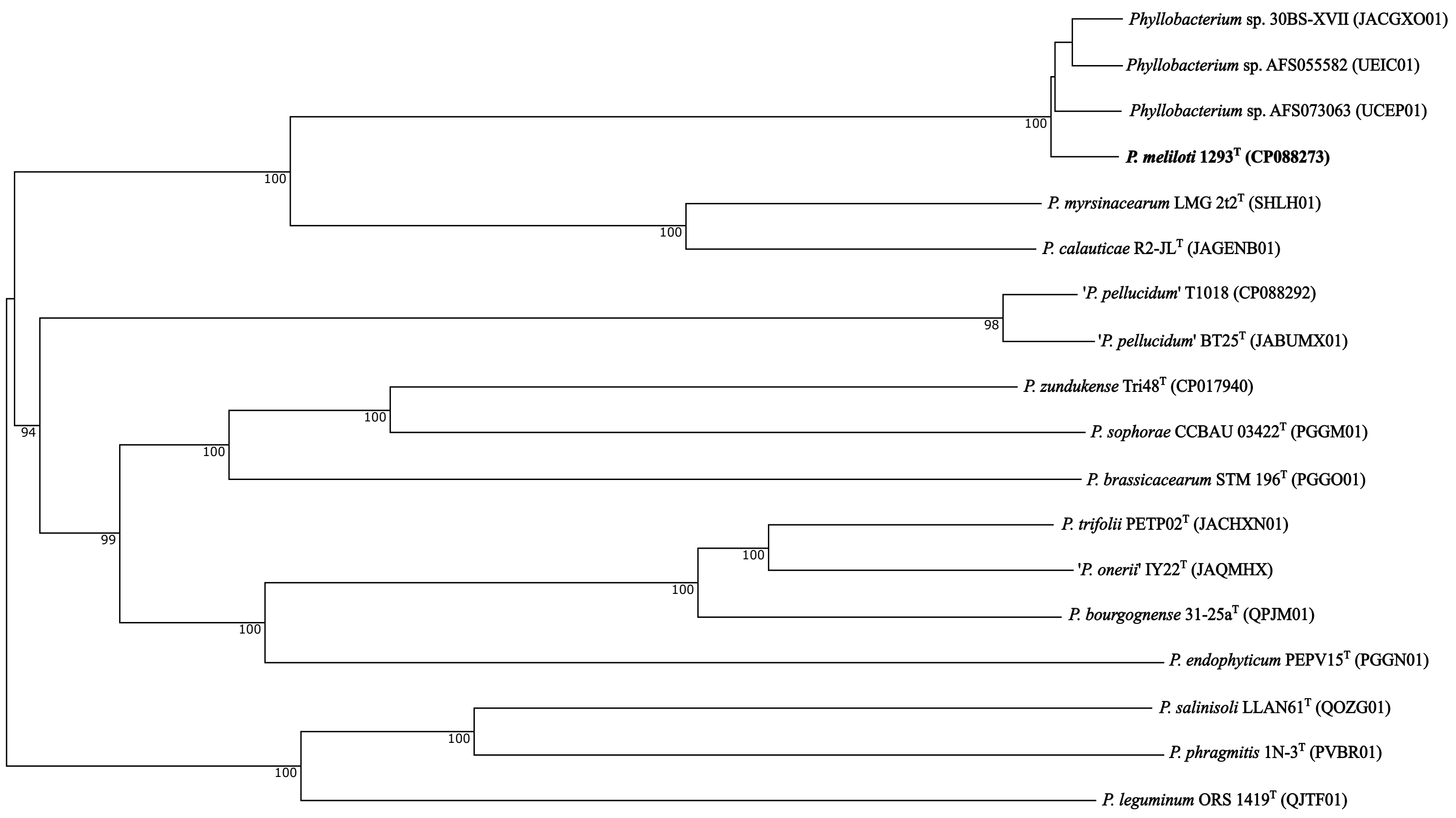


**Supplementary Figure S2.** Phylogenomic tree based on TYGS implementation (Meier-Kolthoff and Göker, 2019) showing *Phyllobacterium meliloti* sp. nov. T1293$^{T}$ and reference taxa of the genus *Phyllobacterium*. The tree was inferred with FastME 2.1.6.1 from Genome Blast Distance Phylogeny (GBDP) distances calculated from genome sequences. Branch lengths are scaled according to GBDP distance formula d5. NCBI sequence accession numbers are given in parentheses. The numbers above branches represent GBDP pseudo-bootstrap support values from 100 replications.

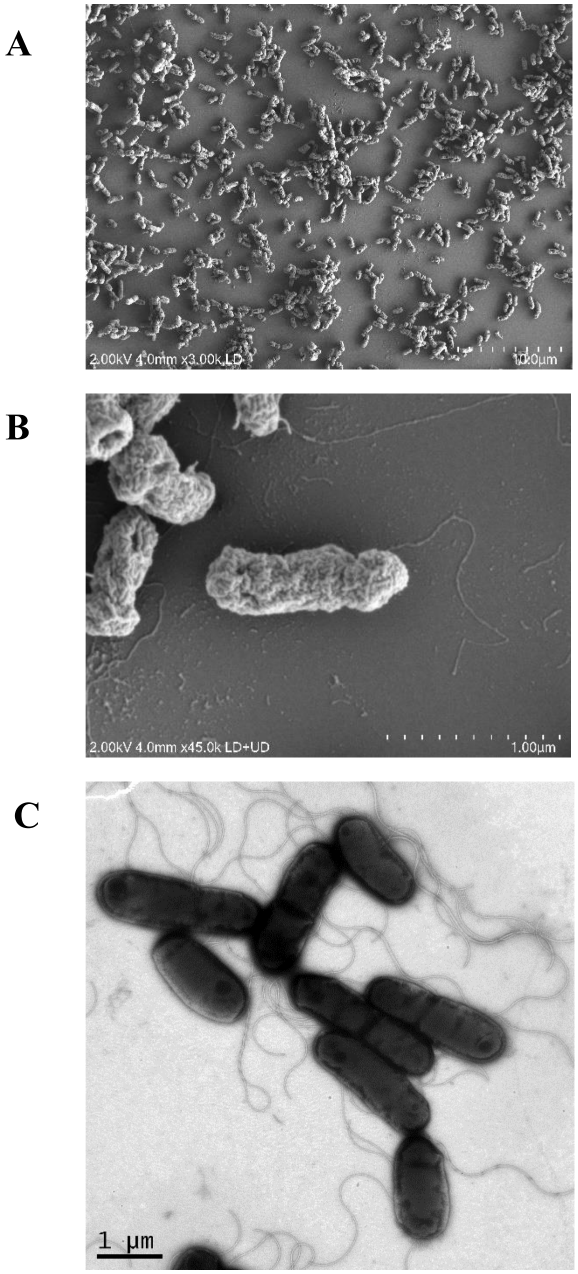


**Supplementary Figure S3.** Scanning (A and B) and transmission (C) electron microscope images showing morphological features of cells of *Phyllobacterium meliloti* T1293$^{T}$.
